## Supplementary Information Adolfi et al. 2020 for "Efficient population modification gene-drive rescue system in the malaria mosquito *Anopheles stephensi*"

This document contains:

- Supplementary Figures 1-6
- Supplementary Tables 1-13

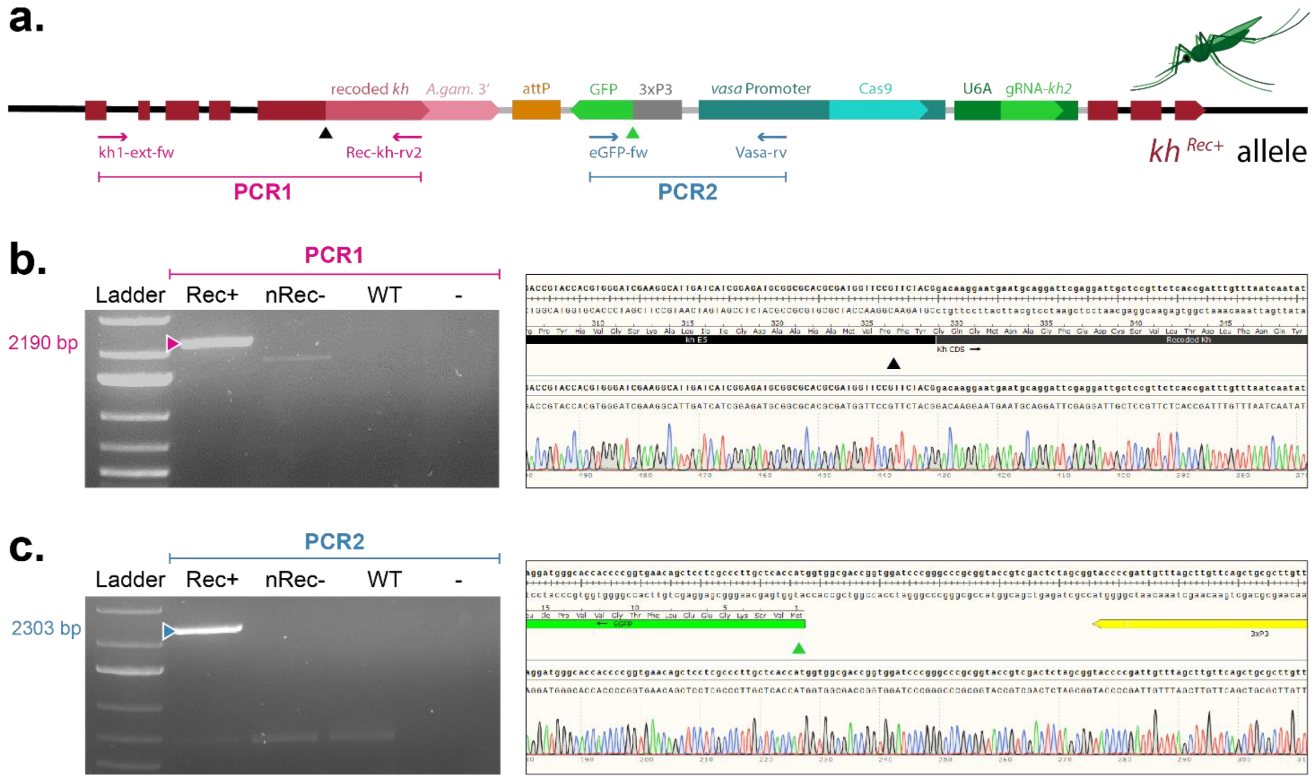

**Supplementary Figure 1.** Confirmation of precise site-specific integration and *kh* recoding in the Rec*kh* gene drive line. **a)** Schematic representation of the integration locus in Rec*kh*, as in Fig. 1b, showing annealing sites of the primers used to verify the integration sites (▲, ▲) of the pRec*kh* recoding element into the nRec line. Recoded *kh* A.gam3': recoded portion of the *kh* cDNA sequence followed by the 3'-end regulatory sequence of the *An. gambiae kh* gene. attP: recombination site for *φC31*-mediated integration. 3xP3-GFP: fluorescent marker driven by an eye specific promoter. *vasa* promoter-Cas9: Cas9 driven by the germline-specific *vasa* promoter. U6A gRNA-kh2: guide RNA targeting *kh* driven by the ubiquitous promoter of the U6A gene. Internal primers Rec-kh-rv2 and eGFP-fw anneal within the pRec*kh* donor element while external primers Kh1-ext-fw and Vasa-rv anneal to sequences outside of the homology arms present in the donor plasmid. **b)** Gene amplification and sequencing of a diagnostic fragment spanning the 5'-end integration site following the cut mediated by gRNA-sw4 (PCR1), which is also the *kh* recoding site. **c)** Gene amplification and sequencing of the 3'-end integration site following the cut mediated by gRNA-sw3 (PCR2). Mosquitoes from the nRec and wild-type (WT) lines were included as controls. Ladder is GeneRuler 1 kb Plus.

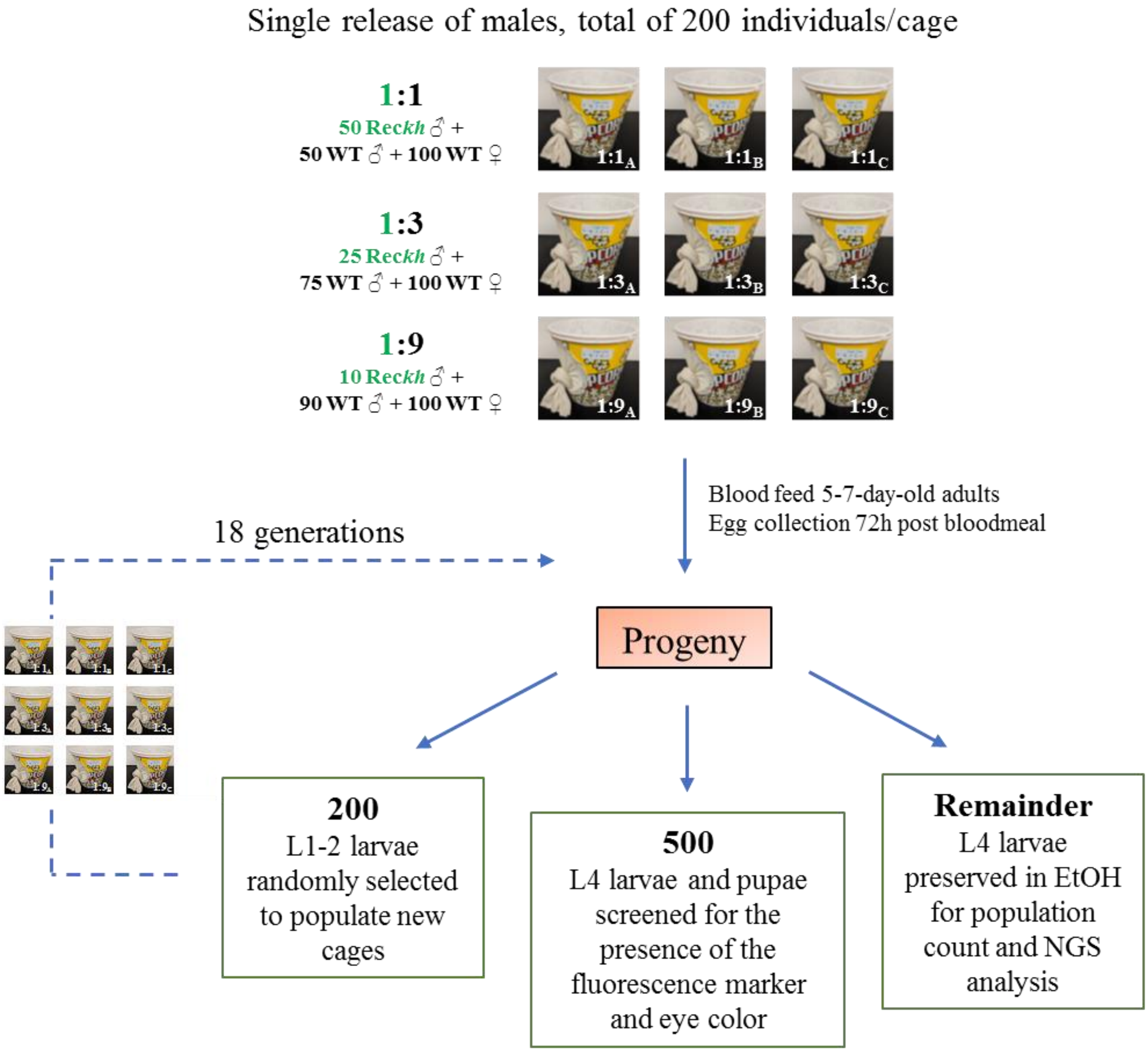

**Supplementary Figure 2.** Schematic representation of the small cage protocol. A total of nine 5,000 $cm^3$ cages were set up with triplicate (A, B and C) initial release ratios of 1:1, 1:3 and 1:9 Rec*kh* drive to wild-type (WT) males. WT females were added to reach a sex ratio of 1:1. A subset of 500 larvae was selected randomly from the progeny obtained from each cage immediately after hatching and screened as L4 larvae and pupae for the presence of the GFP fluorescent marker and the eye color, respectively. A subset of 200 larvae was selected randomly immediately after hatching and reared to adulthood to populate the following set of cages. All individuals from the first generation after release of cages 1:9 were screened and added to seed new cages in proportion to their $GFP^+$ phenotype. The remainder were reared to L4 and stored in ethanol for population counting and sequencing analyses. This protocol was repeated every three weeks for 18 consecutive generations.

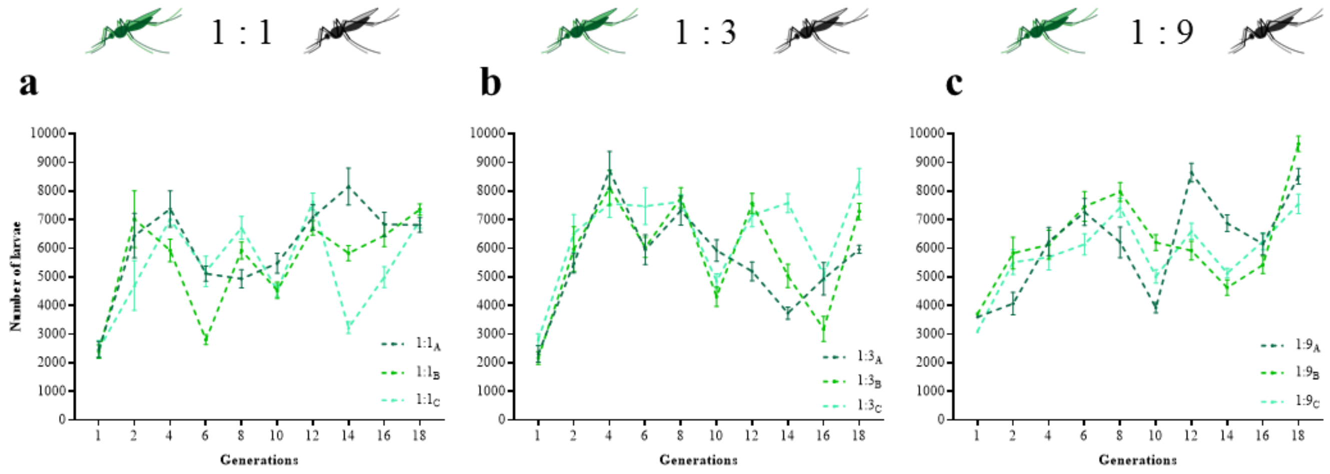

**Supplementary Figure 3.** Population size over time in caged populations seeded with three different release ratios of Rec*kh* to wild-type (WT) males. The total larval population was estimated every two generations in each of the triplicate cages (A, B, and C) seeded with 1:1 (**a**), 1:3 (**b**) and 1:9 (**c**) release ratios of Rec*kh* to WT males by taking 6-9 replicate measurements of randomly-selected L4 larvae.

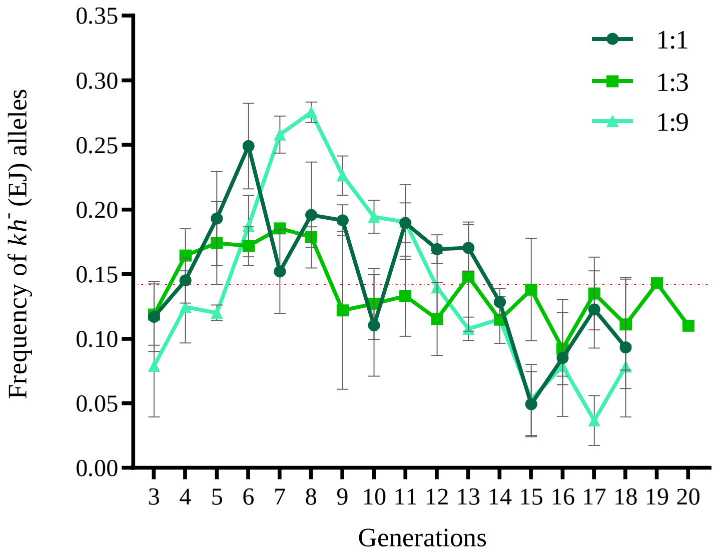

**Supplementary Figure 4.** EJ-induced *kh*⁻ (white) alleles. The frequency of EJ-induced *kh*⁻ alleles over time in all cages was inferred from the number of GFP⁻/white individuals carrying two copies of non-functional mutated alleles using the formula √*N white-eyed individuals / N total mosquitoes* derived from the Hardy-Weinberg equation. Each data point represents the mean with SEM from three replicate cages. Dotted red line represents the average *kh*⁻ allele frequency in all cages during the entire experiment.

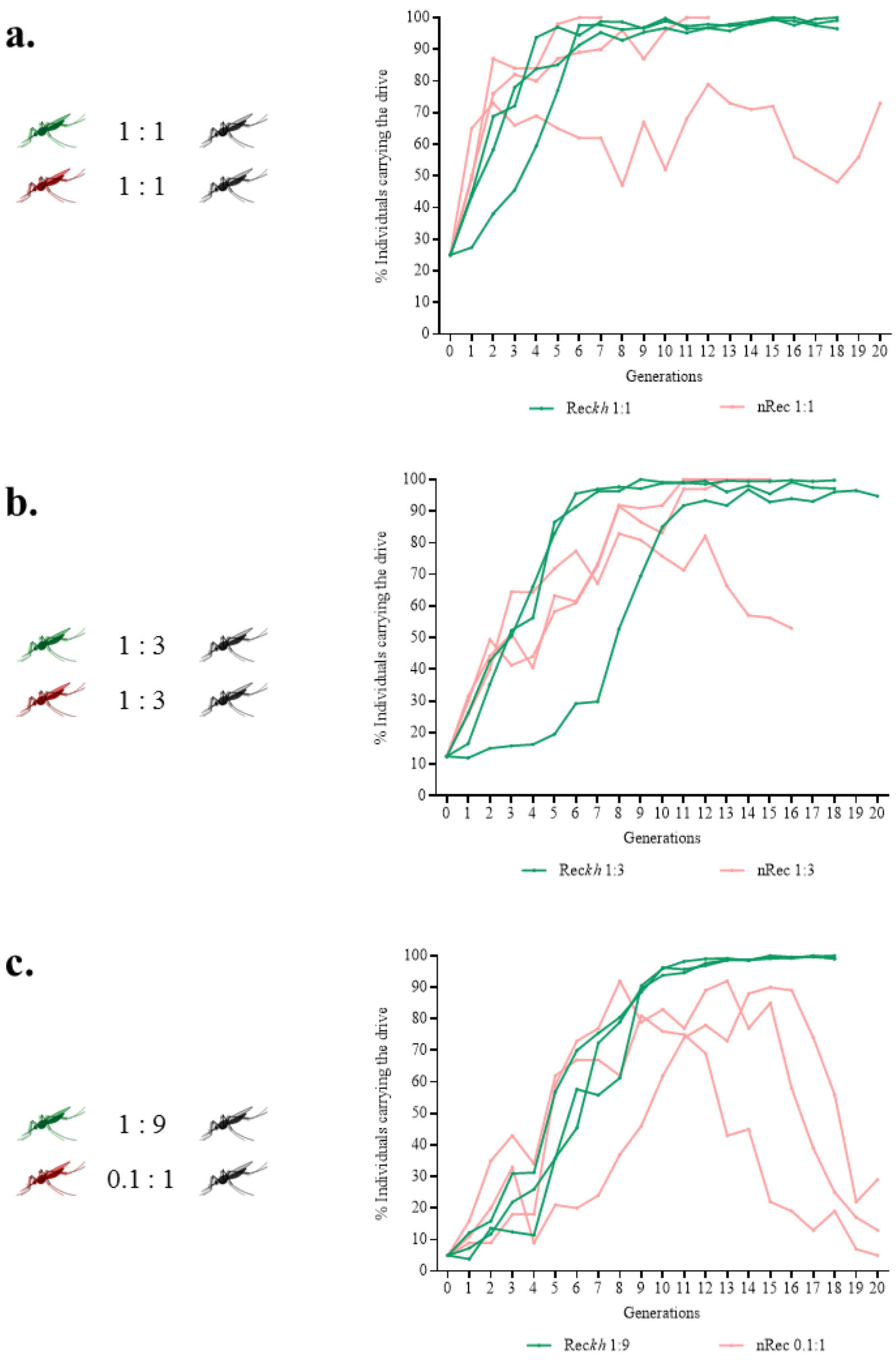

**Supplementary Figure 5.** Comparative long-time dynamics of the Rec*kh* and nRec gene drive lines in small cage laboratory trials. Rec*kh* carries a recoding system that maintains *kh* gene function after insertion, while the insertion of nRec determines the loss-of-function of the *kh* allele. Drive efficiency is measured as the accumulation of the fluorescence marker in three replicate populations of Rec*kh* (green) and nRec (red) *An. stephensi* mosquitoes with initial release ratios of drive to wild-type males of 1:1 (**a**), 1:3 (**b**), and 1:9 (**c**). nRec data are from Pham *et al*.[9].

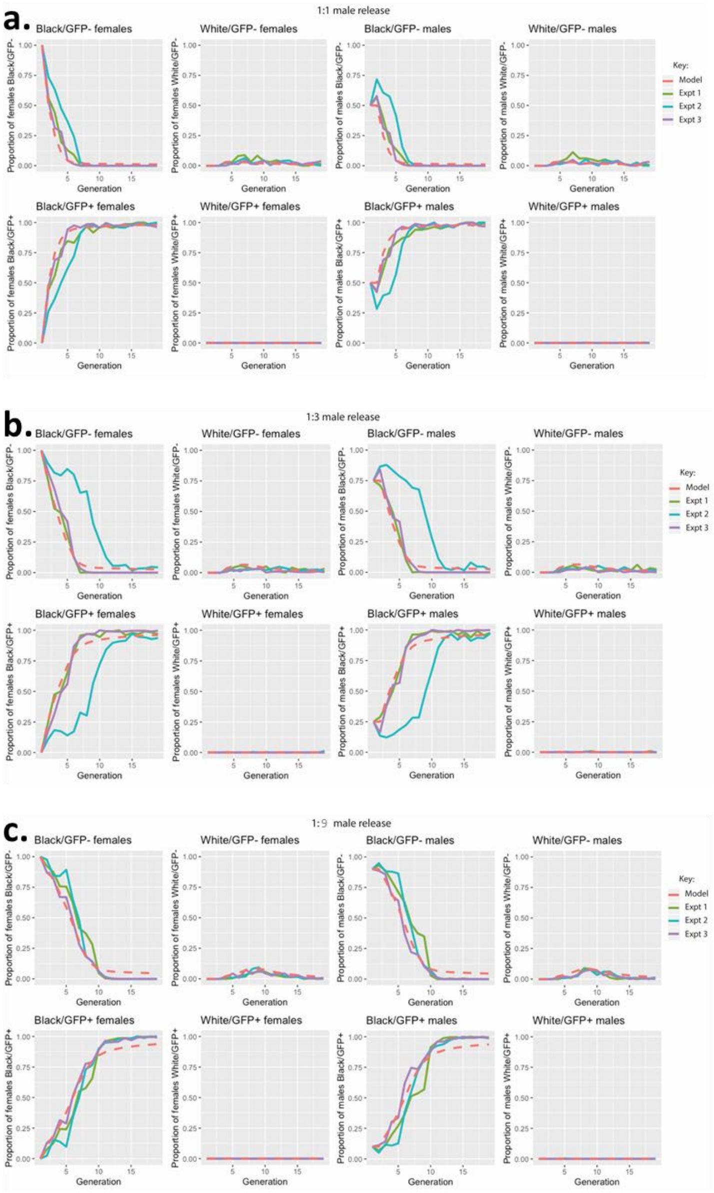

**Supplementary Figure 6.** Observed and model-predicted Rec*kh* gene drive dynamics. Observed (Expt 1-3, solid lines) and predicted (Model, dashed lines) *GFP* and *kh* marker phenotype combinations for 1:1 (**a**), 1:3 (**b**), and 1:9 (**c**) Rec*kh*(HW):wild-type(WW) male releases in 18 non-overlapping generations. *Black*/*GFP*$^{+}$ individuals have at least one copy of the H drive allele (i.e. genotypes HH, HW, HR and HB, where R represents an in-frame, cost-free resistant allele, and B represents an out-of-frame or otherwise costly resistant allele). These individuals spread to near-fixation within ~7 generations (a), 7-12 generations (b), 12 generations (c) of introduction through the inheritance-biasing action of the H allele. *White*/*GFP*$^{-}$ individuals lack both the gene drive construct and a copy of the W or R allele, and hence have genotype BB. B alleles are selected against when present in BB females, but their elimination is slowed due to their viability in heterozygotes and BB males. *Black*/*GFP*$^{-}$ individuals are initially WW, but also include RR, RB, RW and BW genotypes. These genotypes are depleted through the action of the gene drive system but persist at low levels largely due to a small number of R alleles that are generated and persist in the population as a result of their cut-resistant phenotype.

| **Supplementary Table 1. Swap microinjection data.** | | | | | |
|---|---|---|---|---|---|
| **AsMCRkh2 (nRec) embryos** | **$G_0$ larvae** | **$G_0$ adults** | **$G_1$ $GFP^+$ $DsRed^-$ larvae/tot** | **$G_1$ $GFP^+$ $DsRed^-$ adults** | |
| | | | | **From ♂4 founder** | **From ♀4 founder** |
| 504 | 259 | 184 (85 ♂ + 99 ♀) | 96/25,293 | 59 (33 ♂ + 26 ♀) | 13 (7 ♂ + 6 ♀) |

Embryos were obtained from $DsRed^+$ AsMCRkh2 (nRec) females mated to wild-type (WT) males.
$G_0$ individuals were grouped in 27 male founder pools of 2-4 males and 11 female founder pools of 7-10 females and crossed to WT.

**Supplementary Table 2. Drive transmission through Rec*kh* males and females.**

| $G_1$ | $G_2$ | Rep | $G_3$ progeny | | | | | Total (n) | Transmission (% $GFP^+$) | HDR % |
|---|---|---|---|---|---|---|---|---|---|---|
| | | | $GFP^+$ | | $GFP^-$ | | | | | |
| | | | Black | Mosaics | Black | Mosaics | White | | | |
| Hh ♂ | Hh ♂ | A | 149 | 0 | 0 | 0 | 0 | 149 | 100 | 100 |
| | | B | 248 | 0 | 0 | 0 | 0 | 248 | 100 | 100 |
| | | C | 313 | 0 | 0 | 0 | 0 | 313 | 100 | 100 |
| | Hh ♀ | A | 269 | 0 | 0 | 0 | 0 | 269 | 100 | 100 |
| | | B | 258 | 1 | 0 | 0 | 1 | 260 | 99.2 | 98.5 |
| | | C | 329 | 1 | 0 | 0 | 1 | 331 | 99.4 | 98.8 |
| Hh ♀ | Hh ♂ | A | 387 | 0 | 322 | 0 | 0 | 709 | 54.6 | 9.2 |
| | | B | 201 | 0 | 143 | 0 | 0 | 344 | 58.4 | 16.9 |
| | | C | 98 | 0 | 50 | 0 | 0 | 148 | 66.2 | 32.4 |
| | Hh ♀ | A | 217 | 0 | 57 | 1 | 98 | 373 | 58.2 | 16.4 |
| | | B | 228 | 0 | 96 | 0 | 98 | 422 | 54 | 8.1 |
| | | C | 63 | 0 | 24 | 0 | 38 | 125 | 50.4 | 0.9 |

‘H’ is the $kh^{Rec+}$ drive allele; ‘h’ is a non-drive allele.
G1 individuals derived from the cross between HH males and wild-type (hh) females.
All drive individuals were outcrossed to wild-type counterparts *en masse* in three replicate cages (Rep A, B, C).
Transmission % is the proportion of individuals inheriting the drive element ($GFP^+$).
HDR % is the proportion of WT alleles converted into drive alleles by HDR and is calculated using the formula $2(X - 0.5n)/n$ (‘X’ is the number of $GFP^+$ individuals and ‘n’ the total number of mosquito counted).

| **Supplementary Table 3. Reproductive parameters of Rec*kh* and $kh^{-}/kh^{-}$ females.** | | | | | |
|---|---|---|---|---|---|
| | **WT** $kh^{+}/kh^{+}$ | **Rec*kh* homoz** $kh^{Rec+}/kh^{Rec+}$ | **Rec*kh* heteroz** $kh^{Rec+}/kh^{-}$ | **White** $kh^{-}/kh^{-}$ | **One-Way ANOVA** |
| **Feeding success[a]** | 69/75 (92%) | 78/87 (90%) | 96/99 (97%) | 82/94 (87%) | $p = 0.766$ $F (3, 6) = 0.388$ |
| **Survival after blood meal[b]** | 68/69 (99%) | 75/78 (96%) | 96/96 (100%) | 24/82 (29%) | $p = 0.0013$** $F (3, 6) = 21.3$ |
| **Laying females[c]** | 44/59 (75%) | 46/73 (63%) | 48/65 (74%) | 10/24 (42%) | $p = 0.9109$ $F (3, 6) = 0.173$ |
| **Mean No. eggs/female[d] (Fecundity)** | 91 (± 6 SEM) n=31 | 87 (± 5 SEM) n=30 | 89 (± 4 SEM) n=30 | 45 (± 13 SEM) n=10 | $p = 0.0001$*** $F (3, 97) = 7.65$ |
| **Mean No. larvae/female[e] (Fertility)** | 73 (± 5 SEM) n=31 | 75 (± 5 SEM) n=30 | 79 (± 4 SEM) n=30 | 26 (± 8 SEM) n=10 | $p < 0.0001$**** $F (3, 97) = 11.7$ |

[a] Females that fed/total number of females in the cage. 2-3 replicate experiments were performed for each line.
[b] Females that survived a blood meal/total number that fed. 2-3 replicate experiments were performed for each line.
[c] Females that laid eggs/total blood-fed female set up. 2-3 replicate experiments were performed for each line.
[d] Mean (±SEM) number of eggs per laying female.
[e] Mean (±SEM) number of larvae per laying female.
See Supplementary Table 4 for adjusted *p* values after multiple comparisons test.

| **Supplementary Table 4. Adjusted *p* values for parameter in Supplementary Table 3 calculated by applying the Tukey's multiple comparisons test.** | | | |
|---|---|---|---|
| **FEEDING** | $kh^{+}/kh^{+}$ | $kh^{Rec+}/kh^{Rec+}$ | $kh^{Rec+}/kh^{-}$ |
| $kh^{Rec+}/kh^{Rec+}$ | 0.9983 | | |
| $kh^{Rec+}/kh^{-}$ | 0.9482 | 0.8759 | |
| $kh^{-}/kh^{-}$ | 0.9601 | 0.9827 | 0.7211 |
| **SURVIVAL** | $kh^{+}/kh^{+}$ | $kh^{Rec+}/kh^{Rec+}$ | $kh^{Rec+}/kh^{-}$ |
| $kh^{Rec+}/kh^{Rec+}$ | 0.9952 | | |
| $kh^{Rec+}/kh^{-}$ | 0.9996 | 0.9853 | |
| $kh^{-}/kh^{-}$ | 0.0036** | 0.0024** | 0.0033** |
| **LAYING** | $kh^{+}/kh^{+}$ | $kh^{Rec+}/kh^{Rec+}$ | $kh^{Rec+}/kh^{-}$ |
| $kh^{Rec+}/kh^{Rec+}$ | 0.9473 | | |
| $kh^{Rec+}/kh^{-}$ | >0.9999 | 0.9414 | |
| $kh^{-}/kh^{-}$ | 0.9665 | 0.9997 | 0.9619 |
| **EGGS** | $kh^{+}/kh^{+}$ | $kh^{Rec+}/kh^{Rec+}$ | $kh^{Rec+}/kh^{-}$ |
| $kh^{Rec+}/kh^{Rec+}$ | 0.9633 | | |
| $kh^{Rec+}/kh^{-}$ | 0.9949 | 0.9954 | |
| $kh^{-}/kh^{-}$ | <0.0001**** | 0.0004*** | 0.0002*** |
| **LARVAE** | $kh^{+}/kh^{+}$ | $kh^{Rec+}/kh^{Rec+}$ | $kh^{Rec+}/kh^{-}$ |
| $kh^{Rec+}/kh^{Rec+}$ | 0.9949 | | |
| $kh^{Rec+}/kh^{-}$ | 0.7869 | 0.9029 | |
| $kh^{-}/kh^{-}$ | <0.0001**** | <0.0001**** | <0.0001**** |

**Supplementary Table 5. Rec*kh* male contribution to the following generation in the presence of an equal number of wild-type males.**

| Replicate | $GFP^+$ | $GFP^-$ | Tot | % $GFP^+$ | % $GFP^-$ | *p* value* |
|---|---|---|---|---|---|---|
| **Cage A** | 657 | 511 | 1168 | 54.8% | 45.2% | 0.4167 |
| **Cage B** | 639 | 709 | 1348 | 46.7% | 53.3% | 0.6173 |
| **Cage C** | 705 | 769 | 1474 | 46% | 54% | 0.4841 |

Each cage was set up with 75 Rec*kh* homozygous males, 75 wild-type males, and 150 wild-type females.
*Two-tail Binomial Test.

**Supplementary Table 6. Eye phenotypes scored in a subset of ~500 individuals isolated at each generation in the 1:1 cages.**

| Generations | $1{:}1_A$ | | | | | | $1{:}1_B$ | | | | | | $1{:}1_C$ | | | | | |
|---|---|---|---|---|---|---|---|---|---|---|---|---|---|---|---|---|---|---|
| | $GFP^+$ | | | $GFP^-$ | | | $GFP^+$ | | | $GFP^-$ | | | $GFP^+$ | | | $GFP^-$ | | |
| | $kh^+$ Black | $kh^-$ White | $kh^{mos}$ Mos | $kh^+$ Black | $kh^-$ White | $kh^{mos}$ Mos | $kh^+$ Black | $kh^-$ White | $kh^{mos}$ Mos | $kh^+$ Black | $kh^-$ White | $kh^{mos}$ Mos | $kh^+$ Black | $kh^-$ White | $kh^{mos}$ Mos | $kh^+$ Black | $kh^-$ White | $kh^{mos}$ Mos |
| **G1** | 200 | 0 | 0 | 259 | 0 | 0 | 137 | 0 | 0 | 363 | 0 | 0 | 217 | 0 | 0 | 272 | 0 | 0 |
| **G2** | 288 | 0 | 0 | 206 | 0 | 0 | 181 | 0 | 0 | 294 | 0 | 0 | 324 | 0 | 0 | 147 | 0 | 0 |
| **G3** | 368 | 0 | 0 | 87 | 17 | 6 | 203 | 0 | 0 | 235 | 8 | 0 | 351 | 0 | 0 | 134 | 2 | 0 |
| **G4** | 410 | 0 | 0 | 63 | 16 | 1 | 271 | 0 | 0 | 179 | 6 | 0 | 448 | 0 | 0 | 19 | 11 | 1 |
| **G5** | 388 | 0 | 0 | 35 | 33 | 2 | 378 | 0 | 0 | 97 | 16 | 0 | 517 | 0 | 0 | 5 | 11 | 1 |
| **G6** | 452 | 0 | 0 | 1 | 50 | 0 | 455 | 0 | 0 | 16 | 24 | 0 | 445 | 0 | 0 | 0 | 21 | 0 |
| **G7** | 450 | 0 | 0 | 0 | 22 | 0 | 472 | 0 | 0 | 3 | 8 | 0 | 479 | 0 | 0 | 0 | 6 | 0 |
| **G8** | 468 | 0 | 0 | 0 | 36 | 0 | 481 | 0 | 0 | 0 | 19 | 0 | 500 | 1 | 0 | 0 | 7 | 0 |
| **G9** | 456 | 0 | 0 | 0 | 22 | 0 | 471 | 0 | 0 | 0 | 15 | 0 | 457 | 0 | 0 | 0 | 16 | 0 |
| **G10** | 469 | 0 | 0 | 0 | 16 | 0 | 481 | 0 | 0 | 0 | 1 | 0 | 464 | 0 | 0 | 0 | 5 | 0 |
| **G11** | 459 | 0 | 0 | 0 | 23 | 0 | 475 | 0 | 0 | 0 | 17 | 0 | 466 | 0 | 0 | 0 | 13 | 0 |
| **G12** | 486 | 0 | 0 | 0 | 16 | 0 | 466 | 0 | 0 | 0 | 16 | 0 | 501 | 0 | 0 | 0 | 11 | 0 |
| **G13** | 474 | 0 | 0 | 0 | 10 | 0 | 477 | 0 | 0 | 0 | 21 | 0 | 482 | 0 | 0 | 0 | 13 | 0 |
| **G14** | 491 | 0 | 0 | 0 | 6 | 0 | 476 | 0 | 0 | 0 | 8 | 0 | 462 | 0 | 0 | 0 | 10 | 0 |
| **G15** | 486 | 0 | 0 | 0 | 0 | 0 | 472 | 0 | 0 | 0 | 2 | 0 | 436 | 0 | 0 | 0 | 3 | 0 |
| **G16** | 464 | 0 | 0 | 0 | 0 | 0 | 495 | 0 | 0 | 0 | 12 | 0 | 489 | 0 | 0 | 0 | 5 | 0 |
| **G17** | 463 | 0 | 0 | 0 | 9 | 0 | 473 | 0 | 0 | 0 | 2 | 0 | 466 | 1 | 0 | 0 | 12 | 0 |
| **G18** | 459 | 0 | 0 | 0 | 4 | 0 | 475 | 0 | 0 | 0 | 0 | 0 | 467 | 0 | 0 | 0 | 17 | 0 |
| **TOT** | **7731** | **0** | **0** | **651** | **280** | **9** | **7339** | **0** | **0** | **1187** | **175** | **0** | **7971** | **2** | **0** | **577** | **163** | **2** |

**Supplementary Table 7. Eye phenotypes scored in a subset of ~500 individuals isolated at each generation in the 1:3 cages.**

| **Generations** | **1:3$_A$** | | | | | | **1:3$_B$** | | | | | | **1:3$_C$** | | | | | |
|---|---|---|---|---|---|---|---|---|---|---|---|---|---|---|---|---|---|---|
| | **GFP$^+$** | | | **GFP$^-$** | | | **GFP$^+$** | | | **GFP$^-$** | | | **GFP$^+$** | | | **GFP$^-$** | | |
| | ***kh***$^+$ **Black** | ***kh***$^-$ **White** | ***kh***$^{mos}$ **Mos** | ***kh***$^+$ **Black** | ***kh***$^-$ **White** | ***kh***$^{mos}$ **Mos** | ***kh***$^+$ **Black** | ***kh***$^-$ **White** | ***kh***$^{mos}$ **Mos** | ***kh***$^+$ **Black** | ***kh***$^-$ **White** | ***kh***$^{mos}$ **Mos** | ***kh***$^+$ **Black** | ***kh***$^-$ **White** | ***kh***$^{mos}$ **Mos** | ***kh***$^+$ **Black** | ***kh***$^-$ **White** | ***kh***$^{mos}$ **Mos** |
| **G1** | 111 | 0 | 0 | 312 | 0 | 0 | 63 | 0 | 0 | 461 | 0 | 0 | 80 | 0 | 0 | 403 | 0 | 0 |
| **G2** | 207 | 0 | 0 | 279 | 0 | 0 | 73 | 0 | 0 | 413 | 0 | 0 | 168 | 0 | 0 | 309 | 0 | 0 |
| **G3** | 259 | 1 | 0 | 247 | 8 | 4 | 75 | 0 | 0 | 377 | 22 | 9 | 250 | 1 | 0 | 224 | 5 | 2 |
| **G4** | 296 | 1 | 0 | 131 | 21 | 3 | 78 | 0 | 0 | 392 | 10 | 0 | 247 | 0 | 0 | 180 | 12 | 3 |
| **G5** | 409 | 0 | 0 | 56 | 28 | 1 | 101 | 0 | 0 | 402 | 15 | 1 | 449 | 0 | 0 | 62 | 8 | 0 |
| **G6** | 452 | 0 | 0 | 0 | 19 | 0 | 144 | 0 | 0 | 315 | 14 | 3 | 456 | 0 | 0 | 24 | 13 | 0 |
| **G7** | 503 | 2 | 0 | 1 | 15 | 0 | 145 | 2 | 0 | 330 | 16 | 1 | 456 | 1 | 0 | 2 | 16 | 0 |
| **G8** | 470 | 2 | 0 | 0 | 11 | 0 | 257 | 0 | 0 | 214 | 16 | 0 | 476 | 0 | 0 | 0 | 18 | 0 |
| **G9** | 473 | 1 | 0 | 0 | 14 | 0 | 342 | 0 | 0 | 132 | 18 | 0 | 495 | 0 | 0 | 0 | 0 | 0 |
| **G10** | 496 | 0 | 0 | 0 | 6 | 0 | 390 | 0 | 0 | 54 | 15 | 0 | 481 | 0 | 0 | 0 | 4 | 0 |
| **G11** | 477 | 0 | 0 | 0 | 5 | 0 | 435 | 0 | 0 | 21 | 18 | 0 | 478 | 0 | 0 | 0 | 5 | 0 |
| **G12** | 492 | 0 | 0 | 0 | 2 | 0 | 426 | 0 | 0 | 18 | 12 | 0 | 479 | 0 | 0 | 0 | 7 | 0 |
| **G13** | 466 | 0 | 0 | 0 | 19 | 0 | 445 | 0 | 0 | 24 | 16 | 0 | 482 | 0 | 0 | 0 | 2 | 0 |
| **G14** | 463 | 0 | 0 | 0 | 9 | 0 | 479 | 0 | 0 | 8 | 8 | 0 | 479 | 0 | 0 | 0 | 3 | 0 |
| **G15** | 465 | 0 | 0 | 0 | 22 | 0 | 431 | 0 | 0 | 26 | 7 | 0 | 484 | 0 | 0 | 0 | 3 | 0 |
| **G16** | 493 | 0 | 0 | 0 | 4 | 0 | 468 | 0 | 0 | 20 | 10 | 0 | 489 | 0 | 0 | 0 | 1 | 0 |
| **G17** | 453 | 2 | 0 | 0 | 12 | 0 | 443 | 0 | 0 | 22 | 11 | 0 | 467 | 0 | 0 | 0 | 3 | 0 |
| **G18** | 488 | 0 | 0 | 0 | 14 | 0 | 464 | 3 | 0 | 15 | 4 | 0 | 476 | 0 | 0 | 0 | 1 | 0 |
| **G19** | - | - | - | - | - | - | 471 | 0 | 0 | 7 | 10 | 0 | - | - | - | - | - | - |
| **G20** | - | - | - | - | - | - | 473 | 0 | 0 | 20 | 6 | 0 | - | - | - | - | - | - |
| **TOT** | **7473** | **9** | **0** | **1026** | **209** | **8** | **6203** | **5** | **0** | **3271** | **228** | **14** | **7392** | **2** | **0** | **1204** | **101** | **5** |

**Supplementary Table 8. Eye phenotypes scored in a subset of ~500 individuals isolated at each generation in the 1:9 cages.**

| Generations | $1:9_A$ | | | | | | $1:9_B$ | | | | | | $1:9_C$ | | | | | |
|---|---|---|---|---|---|---|---|---|---|---|---|---|---|---|---|---|---|---|
| | $GFP^+$ | | | $GFP^-$ | | | $GFP^+$ | | | $GFP^-$ | | | $GFP^+$ | | | $GFP^-$ | | |
| | $kh^+$ Black | $kh^-$ White | $kh^{mos}$ Mos | $kh^+$ Black | $kh^-$ White | $kh^{mos}$ Mos | $kh^+$ Black | $kh^-$ White | $kh^{mos}$ Mos | $kh^+$ Black | $kh^-$ White | $kh^{mos}$ Mos | $kh^+$ Black | $kh^-$ White | $kh^{mos}$ Mos | $kh^+$ Black | $kh^-$ White | $kh^{mos}$ Mos |
| **G1** | 36 | 0 | 0 | 457 | 0 | 0 | 19 | 0 | 0 | 479 | 0 | 0 | 62 | 0 | 0 | 447 | 0 | 0 |
| **G2** | 56 | 0 | 0 | 419 | 0 | 0 | 69 | 0 | 0 | 437 | 0 | 0 | 79 | 0 | 0 | 418 | 0 | 0 |
| **G3** | 97 | 0 | 0 | 347 | 0 | 0 | 65 | 0 | 0 | 449 | 10 | 2 | 145 | 0 | 0 | 315 | 9 | 3 |
| **G4** | 118 | 1 | 0 | 334 | 6 | 2 | 56 | 0 | 0 | 428 | 6 | 2 | 145 | 0 | 0 | 300 | 19 | 4 |
| **G5** | 170 | 0 | 0 | 297 | 8 | 0 | 171 | 0 | 0 | 297 | 8 | 1 | 287 | 0 | 0 | 210 | 8 | 2 |
| **G6** | 266 | 0 | 1 | 214 | 18 | 1 | 242 | 0 | 0 | 210 | 11 | 1 | 373 | 0 | 0 | 128 | 32 | 4 |
| **G7** | 275 | 0 | 0 | 182 | 36 | 2 | 342 | 0 | 0 | 92 | 39 | 2 | 354 | 0 | 0 | 86 | 29 | 4 |
| **G8** | 297 | 0 | 0 | 154 | 34 | 0 | 372 | 0 | 0 | 59 | 40 | 0 | 376 | 0 | 0 | 57 | 34 | 0 |
| **G9** | 455 | 0 | 0 | 19 | 29 | 0 | 445 | 0 | 0 | 31 | 20 | 1 | 436 | 0 | 0 | 27 | 30 | 1 |
| **G10** | 430 | 0 | 0 | 1 | 17 | 0 | 441 | 0 | 0 | 7 | 22 | 0 | 453 | 0 | 0 | 4 | 14 | 0 |
| **G11** | 487 | 0 | 0 | 0 | 9 | 0 | 471 | 0 | 0 | 0 | 27 | 0 | 489 | 0 | 0 | 1 | 21 | 0 |
| **G12** | 506 | 0 | 0 | 1 | 4 | 0 | 481 | 0 | 0 | 0 | 12 | 0 | 473 | 0 | 0 | 0 | 15 | 0 |
| **G13** | 494 | 0 | 0 | 0 | 4 | 0 | 452 | 0 | 0 | 0 | 6 | 0 | 489 | 0 | 0 | 0 | 7 | 0 |
| **G14** | 461 | 0 | 0 | 1 | 6 | 0 | 460 | 0 | 0 | 0 | 6 | 0 | 478 | 0 | 0 | 0 | 7 | 0 |
| **G15** | 472 | 0 | 0 | 0 | 0 | 0 | 466 | 0 | 0 | 0 | 2 | 0 | 462 | 0 | 0 | 0 | 4 | 0 |
| **G16** | 486 | 0 | 0 | 0 | 2 | 0 | 481 | 0 | 0 | 0 | 4 | 0 | 445 | 0 | 0 | 0 | 3 | 0 |
| **G17** | 476 | 0 | 0 | 0 | 2 | 0 | 483 | 0 | 0 | 0 | 1 | 0 | 506 | 0 | 0 | 0 | 0 | 0 |
| **G18** | 489 | 0 | 0 | 0 | 4 | 0 | 485 | 1 | 0 | 0 | 0 | 0 | 491 | 0 | 0 | 0 | 5 | 0 |
| **TOT** | **6071** | **1** | **1** | **2426** | **179** | **5** | **6001** | **1** | **0** | **2489** | **214** | **9** | **6543** | **0** | **0** | **1993** | **237** | **18** |

**Supplementary Table 9. Sequences of non-drive alleles in non-drive white-eyed (GFP[−]/*kh*[−]) individuals from all cages at generation G3.**

C: cage; I: individual mosquito; WT: wild-type; gRNA; PAM; mutation; F: frame; FS: frameshift.; IF: in frame; HOM: homozygous.

| C | I | Allele 1 | Allele 2 | F |
|---|---|---|---|---|
| | WT | CACGCGATGGTTCCGTTCTACGGGCAGGGCATGAACGCGGG | CACGCGATGGTTCCGTTCTACGGGCAGGGCATGAACGCGGG | |
| 1:1A | 1 | CACGCGATGGTTCCGTTCTACGCAGGCAGGGCATGAACGCGGG | CACGCGATGGTTCCGTTTATGGAT--GGGCATGAACGCGGG | FS/FS |
| | 2 | CACGCGATGGTTCCGTTCTGCACGAAAAGGGCAGGGCATGAACGCGGG | CACGCGATGGTTCCGTTCTACAGGCGCTGGATCAAGGCAGGGCATGAACGCGGG | FS/FS |
| | 3 | CACGCGATGGTTCCGTTCTAC-GGCAGGGCATGAACGCGGG | CACGCGATGGTTCCGTTCTACAGGGCAGGGCATGAACGCGGG | FS/FS |
|  | 4 | CACGCGATGGTTCCGTTCTACAGGCAGGGCATGAACGCGGG | CACGCGATGGTTCCGTTCTAC----AGGGCATGAACGCGGG | FS/FS |
|  | 5 | CACGCGATGGTTCCGTTCAT----CAGGGCATGAACGCGGG | CACGCGATGGTTCCGTTCTA--GGCAGGGCATGAACGCGGG | FS/FS |
| | 6 | CACGCGATGGTTCCGTTCTACCGGGCAGGGCATGAACGCGGG | CACGCGATGGTTCCGTTCTAAATCGCAGGCAGGGCATGAACGCGGG | FS/FS |
| | 7 | CACGCGATGGTTCCGTTCTACCAGGGCAGGGCATGAACGCGGG | CACGCGATGGTTCCGTTCTACAGTCGCACGGCAGGGCATGAACGCGGG | FS/FS |
| | 8 | CACGCGATGGTTCCGTTCTAC-GGCAGGGCATGAACGCGGG | CACGCGATGGTTCCGTTCGAT-GGCAGGGCATGAACGCGGG | FS/FS |
| | 9 | CACGCGATGGTTCCGT---ACGGGCAGGGCATGAACGCGGG | CACGCGATGGTTCCGTTTCCCCGGCAGGGCATGAACGCGGG | IF/IF |
| 1:1B | 1 | CACGCGATGGTTCCGTTCTACG--CAGGGCATGAACGCGGG | CACGCGATGGTTCCGTTCTACAGGCAGGGCATGAACGCGGG | FS/IF |
| | 2 | CACGCGATGGTTCCGTTCTACG--CAGGGCATGAACGCGGG | CACGCGATGGTTCCGTTCTACGATGGCAGGGCATGAACGCGGG | FS/FS |
| | 3 | CACGCGATGGTTCCGTTCT---GGCAGGGCATGAACGCGGG | CACGCGATGGTTCCGTTCCACG--CAGGGCATGAACGCGGG | IF/FS |
| | 4 | CACGCGATGGTTCCGTTCTACG--CAGGGCATGAACGCGGG | CACGCGATGGTTCCGTTCTACGGGACCAGGCAGGGCATGAACGCGGG | FS/IF |
|  | 5 | CACGCGATGGTTCCG--------GCAGGGCATGAACGCGGG | CACGCGATGGTTCCGTTCC---GGCAGGGCATGAACGCGGG | FS/IF |
| | 6 | CACGCGATGGTTCCGTTCTACGCG--GGGCATGAACGCGGG | CACGCGATGGTTCCGTTCTACAGGGAACCATCAAAGGCACGTCAAGAACGCGGG | FS/FS |
| | 7 | CACGCGATGGTTCCGTTCTACGCAGGGCAGGGCATGAACGCGGG | CACGCGATGGTTCCGTTCGCGGCAGGGCAGGGCATGAACGCGGG | IF/IF |
| | 8 | CACGCGATGGTTCCGTTCA---GGCAGGGCATGAACGCGGG | CACGCGATGGTTCCGTTCTTGCACGCGTTGGCAGGGCATGAACGCGGG | FS/FS |
| 1:1C | 1 | CACGCGATGGTTCCGTTCT---GGCAGGGCATGAACGCGGG | CACGCGATGGTTCCGTTCCACAGGGGCAGGGCATGAACGCGGG | IF/FS |
|  | 2 | CACGCGATGGTTCCGTTCTACGGGCATGGCAGGGCATGAACGCGGG | CACGCGATGGTTCCG------GCGGGCATTGCATGAACGCGGG | FS/FS |
| 1:3A | 1 | CACGCGATGGTTCCGTT-----GGCAGGGCATGAACGCGGG | CACGCGATGGTTCCGTTCTACGCAGGGCAGGGCATGAACGCGGG | FS/FS |

| | | | | |
|---|---|---|---|---|
|  | 2 | CACGCGATGGTTCCGTTCTAC--GCAGGGCATGAACGCGGG | CACGCGATGGTTCCGAT-----GGCAGGGCATGAACGCGGG | FS/FS |
| 1:3B | 1 | CACGCGATGGTTC-----TACG--CAGGGCATGAACGCGGG | CACGCGATGGTTC-----TACG--CAGGGCATGAACGCGGG | FS HOM |
|  | 2 | CACGCGATGGTTCCGTTCTAC--GCAGGGCATGAACGCGGG | -------------------------AGGGCATGAACGCGGG | FS/FS |
| | 3 | CACGCGATGGTTCCGTTCTACAGGGCAGGGCATGAACGCGGG | CACGCGATGGTTCCGTTCTACGCGATGGTTCCGGCAGGGCATGAACGCGGG | FS/FS |
| | 4 | CACGCGATGGTTCCGTTCTAC--GCAGGGCATGAACGCGGG | CACGCGATGGTTCCGTTCTACAGGGCAGGGCATGAACGCGGG | FS/FS |
| | 5 | CACGCGATGGTTCCGTTCTACG--CAGGGCATGAACGCGGG | CACGCGATGGTTCCGTTCTACGCGATGGCAGGGCATGAACGCGGG | FS/FS |
| | 6 | CACGCGATGGTTCCGTTCACGCGGGCAGGGCATGAACGCGGG | CACGCGATGGTTCCGTTCTACAGGCAGGGCATGAACGCGGG | FS/IF |
| | 7 | CACGCGATGGTTCCGTTCGCCAGGGCAGGGCATGAACGCGGG | CACGCGATGGTTCCGTTCTAGTTCCGGCAGGGCATGAACGCGGG | FS/FS |
| | 8 | CACGCGATGGTTCCGTTCTACGCCGGCAGGGCATGAACGCGGG | CACGCGATGGTTCCGTTCCGTT-GCAGGGCATGAACGCGGG | FS/FS |
| | 9 | CACGCGATGGTTCCGTTCTA--GGCAGGGCATGAACGCGGG | CACGCGATGGTTCCGTTCTACCGCGCGATGGTTCCGTGGTTCCGGCAGGGCATGAACGCGGG | FS/FS |
| 1:3C | 1 | CACGCGATGGTTCCGTTCTAC----AGGGCATGAACGCGGG | CACGCGATGGTTCCGTTCTACATGAACGGCAGGGCATGAACGCGGG | FS/FS |
| 1:9B | 1 | CACGCGATGGTTCCGTTCTACGGGCA-----TGAACGCGGG | CACGCGATGGTTCCGTTCTACAG-CAGGGCATGAACGCGGG | FS/FS |
| | 2 | CACGCGATGGTTCCGTTCTACTCCGGCAGGGCATGAACGCGGG | CACGCGATGGTTCCGTTCTACGGAACGTTCACGGCAGGGCATGAACGCGGG | FS/FS |
| | 3 | CACGCGATGGTTCCGTTCTACG-GCAGGGCATGAACGCGGG | CACGCGATGGTTCCGTTCTACG-GCAGGGCATGAACGCGGG | FS HOM |
| 1:9C | 1 | CACGCGATGGTTCCGTTCTACAAGGGCAGGGCATGAACGCGGG | CACGCGATGGTTCCGTTCCGCGTGAACGGCGGCCTGGTTCATGGTATGACATGAACGCGGG | FS/FS |
| | 2 | CACGCGATGGTTCCGTTCC---GGCAGGGCATGAACGCGGG | CACGCGATGGTTCCGTTCCGCGATGGCAGGGCATGAACGCGGG | IF/FS |
|  | 3 | CACGCGATGGTTCCGAT-----GGCAGGGCATGAACGCGGG | CACGCGATGGTTCCGAT-----GGCAGGGCATGAACGCGGG | FS HOM |
| | 4 | CACGCGATGGTTCCGTTCTAC-GGCAGGGCATGAACGCGGG | CACGCGATGGTTCCGTTCTACTCGAAATCACGCGATGGTTCCGTTCTAAGGCAGGGCATGAACGCGGG | FS/FS |
|  | 5 | CACGCGATGGTTC-----TACG--CAGGGCATGAACGCGGG | CACGCGATGG------TTCTACGGCAGGGCATGAACGCGGG | FS/FS |

**Supplementary Table 10. Sequences of drive and non-drive alleles in drive white-eyed (GFP$^{+}$/*kh*$^{-}$) individuals.**

C: Cage; G: generation; I: individual mosquito; WT: wild-type; gRNA; PAM; mutation; Recoded-*kh*.

| C | G | I | Non-drive Allele | Rec*kh* Drive Allele |
|---|---|---|---|---|
| | | WT | CACGCGATGGTTCCGTTCTACGGGCAGGGCATGAACGCGGG | CACGCGATGGTTCCGTTCTACGGACAAGGAATGAATGCAGGATTC |
| 1:1$_C$ | G8 | 1 | CACGCGATGGTTCCGTTCTACATGAACGCGGCAGGGCATGAACGCGGG | CACGCGATGGTTCCGTTCCGATGGTTCCGGACAAGGAATGAATGCAGGATTC |
| 1:3$_A$ | G7 | 2 | CACGCGATGGTTCCGTTCTACAGGGCAGGGCATGAACGCGGG | CACGCGATGGTTCCGTTCCGATGGTTCCGGACAAGGAATGAATGCAGGATTC |
| 1:3$_A$ | G7 | 3 | CACGCGATGGTTCCGTTCTACGC-CAGGGTATGAACGCGGG | CACGCGATGGTTCCGTTC----GACAAGGAATGAATGCAGGATTC |
| 1:3$_A$ | G8 | 4 | CACGCGATGGTTCCGTTCTACGC-CAGGGTATGAACGCGGG | CACGCGATGGTTCCGTTC----GACAAGGAATGAATGCAGGATTC |
| 1:3$_A$ | G8 | 5 | CACGCGATGGTTCCGTTCTACGG-CAGGGCATGAACGCGGG | CACGCGATGGTTCCGTTCTACAAGGAAGGGACAAGGAATGAATGCAGGATTC |
| 1:9$_A$ | G4 | 6 | CACGCGATGGTTCCGTTCC---GGCAGGGCATGAACGCGGG | CACGCGATGGTTCCGTTCTGGACAAGGACAAGGAATGAATGCAGGATTC |

| **Supplementary Table 11. Amplicon sequencing of non-drive alleles in pooled individuals from cage 1:3B at generations G0, G8, and G14.** | | |
|---|---|---|
| gRNA and PAM in the wild-type allele; Mutation. | | |
| **Cage 1:3B Generation G0** | | |
| **Reads** | **Sequence** | **Relative abundance (%)** |
| 315911 | GATGGTTCCGTTCTACGGGCAGGGCATGAACGCG | 97.65 |
| 938 | GATGGTTCCGTTCTACGGGCAGGACATGAACGCG | 0.29 |
| 730 | GATGGT-----------GGCAGGGCATGAACGCG | 0.23 |
| 600 | GATGGTTCCGTTCTACGGACAGGGCATGAACGCG | 0.19 |
| 583 | GATGG------------GGCAGGGCATGAACGCG | 0.18 |
| 500 | GATGGTTCCGTT-----GGCAGGGCATGAACGCG | 0.15 |
| 391 | GATGGTTCCGT------GGCAGGGCATGAACGCG | 0.12 |
| 357 | GATGGTT----------GGCAGGGCATGAACGCG | 0.11 |
| 343 | GATGGTTCC--------GGCAGGGCATGAACGCG | 0.11 |
| 325 | GATGGTTCCGTTCTCCGGGCAGGGCATGAACGCG | 0.10 |
| 235 | GATGGTTCCGTTCC---GGCAGGGCATGAACGCG | 0.07 |
| 229 | GATGGTTCCGTTCTA--GGCAGGGCATGAACGCG | 0.07 |
| 194 | GATG-------------GGCAGGGCATGAACGCG | 0.06 |
| 190 | GATGGTTCCG-------GGCAGGGCATGAACGCG | 0.06 |
| 183 | GATGGTTCCGTTCT---GGCAGGGCATGAACGCG | 0.06 |
| 161 | GATGGTTCCGTTCTACGAGCAGGGCATGAACGCG | 0.05 |
| 158 | GATGGTTCCGTTC----GGCAGGGCATGAACGCG | 0.05 |
| 156 | GATGG--------------CAGGGCATGAACGCG | 0.05 |
| 139 | GATGGTTCCGTTCTACG-GCAGGGCATGAACGCG | 0.04 |
| 136 | GATGGTTCCGTTC-------AGGGCATGAACGCG | 0.04 |
| 129 | GATGGTTC---------GGCAGGGCATGAACGCG | 0.04 |
| 129 | GATGGTTCCGTTCTATGGGCAGGGCATGAACGCG | 0.04 |
| 126 | GATGGTTCCGTTCTACGGGCAGGGCATGGACGCG | 0.04 |

| | | |
|---|---|---|
| 122 | AATGGTTCCGTTCTACGGGCAGGGCATGAACGCG | 0.04 |
| 115 | GATGGTTCCGTTCTACGGGCAGTGCATGAACGCG | 0.04 |
| 111 | GATGGTTCCGTTCTACGGGCAGGGCATGAACACG | 0.03 |
| 103 | GATGGTTCCGTTCTAC----AGGGCATGAACGCG | 0.03 |
| 103 | GATGGTTCCGTTC--ATGGCAGGGCATGAACGCG | 0.03 |
| 101 | GATGGTTCCGTTCTACGGGCAGGGCATGAACGCT | 0.03 |
| **323498** | | |
| **Cage 1:3B Generation G8** | | |
| **Reads** | **Sequence** | **Relative abundance (%)** |
| 175157 | GATGGTTCCGTTCTACGGGCAGGGCATGAACGCG | 83.28 |
| 6438 | GATGGTTCCGTTCTACAGGGCAGGGCATGAACGCG | 3.06 |
| 4558 | GATGGTTCCGTTCTACTAAACACGCGTTGCCATGAACGCGGTTCTACTAAACAGGCAGGGCATGAACGCG | 2.17 |
| 4382 | GATGGTTCCGTTCTACGCATGAACGCAGGGCATGAACGCG | 2.08 |
| 4137 | GATGGTTCCGTTCCGATGGCAGGGCATGAACGCG | 1.97 |
| 4123 | GATGGTTCCGTTCTACAACGCAACGTTCTACAACGGGGCAGGGCATGAACGCG | 1.96 |
| 4042 | GATGGTTCCGTTCC---GGCAGGGCATGAACGCG | 1.92 |
| 2124 | GATGGTTCCGTTCTACGGGGCAGGCATGAACGCG | 1.01 |
| 2085 | GATGGTTACGTTCTACATGGCAGGGCATGAACGCG | 0.99 |
| 2083 | GATGGTTCCGTTCTACGTTCGCGGCAGGGCATGAACGCG | 0.99 |
| 478 | GATGGTTCCGTTCTACGGACAGGGCATGAACGCG | 0.23 |
| 473 | GATGGTTCCGTTCTACGGGCAGGACATGAACGCG | 0.22 |
| 241 | GATGGTTCCGTTCTCCGGGCAGGGCATGAACGCG | 0.11 |
| **210321** | | |
| **Cage 1:3B Generation G14** | | |
| **Reads** | **Sequence** | **Relative abundance (%)** |
| 147513 | GATGGTTCCGTTCTACAACGCAACGTTCTACAACGGGGCAGGGCATGAACGCG | 48.16 |
| 77514 | GATGGTTCCGTTCTACGT----------------- | 25.31 |
| 35534 | GATGGTTCCGTTCTACGGGGCAGGCATGAACGCG | 11.60 |
| 22641 | GATGGTTCCGTTCTACGGAACCGGCAGGCATGAACGCG | 7.39 |

| | | |
|---|---|---|
| 14200 | GATGGTTCCGTTCTACCAGCGCAGGGCAGGGCATGAACGCG | 4.64 |
| 4036 | GATGGTTCCGTTCATGT----GGGCATGAACGCG | 1.32 |
| 1091 | GATGGTTCCGTTCCCGCGTGGGCAGGCATGAACGCG | 0.36 |
| 796 | GATGG---------------CAGGCATGAACGCG | 0.26 |
| 661 | GATGGTTCCGTTCTACAACGCAACGTTCTACAACTGGGCAGGGCATGAACGCG | 0.22 |
| 623 | GATGGTTCCGTTCTACAACGCACCGTTCTACAACGGGGCAGGGCATGAACGCG | 0.20 |
| 593 | G-----------------GGCAGGCATGAACGCG | 0.19 |
| 455 | GATGGTTCCGTTCTACAACGCAACGTTCTACACCGGGGCAGGGCATGAACGCG | 0.15 |
| 236 | GATGGTTCCGTTCTACGGGCAGGGCATGAACGCG | 0.08 |
| 151 | GATGGTTCCGTTCTACAACACAACGTTCTACAACGGGGCAGGGCATGAACGCG | 0.05 |
| 134 | GATGGTTCCGTTCTACAACGCAACGTTATACAACGGGGCAGGGCATGAACGCG | 0.04 |
| 107 | GATGGTTCCGTTCTACGGGGCAGACATGAACGCG | 0.03 |
| **306285** | | |

**Supplementary Table 12. Sequences of non-drive alleles in single non-drive black-eyed (GFP⁻/*kh*⁺) individuals from cage 1:3B at generation G16.**

I: individual mosquito; WT: wild-type; gRNA; PAM; mutation; F: frame; IF: in frame; FS: frameshift; HOM: homozygous; AAC: amino acid change in IF allele.

| I | Allele 1 | Allele 2 | F | AAC |
|---|---|---|---|---|
| WT | CACGCGATGGTTCCGTTCTACGGGCAGGGCATGAACGCGGG | CACGCGATGGTTCCGTTCTACGGGCAGGGCATGAACGCGGGCTTTGAAGACTGTAGC | | |
| 1 | CACGCGATGGTTCCGTTCTACGGGGCAGGCATGAACGCGGG | CACGCGATGGTTCCGTTCTACGGGGCAGGCATGAACGCGGGCTTTGAAGACTGTAGC | IF/IF HOM | Q330A |
| 2 | CACGCGATGGTTCCGTTCTACGGGGCAGGCATGAACGCGGG | CACGCGATGGTTCCGTTCTACGGGGCAGGCATGAACGCGGGCTTTGAAGACTGTAGC | IF/IF HOM | Q330A |
| 3 | CACGCGATGGTTCCGTTCTACGGGGCAGGCATGAACGCGGG | CACGCGATGGTTCCGTTCTACGGGGCAGGCATGAACGCGGGCTTTGAAGACTGTAGC | IF/IF HOM | Q330A |
| 4 | CACGCGATGGTTCCGTTCTACGGGGCAGGCATGAACGCGGG | CACGCGATGGTTCCGTTCTACGGGGCAGGCATGAACGCGGGCTTTGAAGACTGTAGC | IF/IF HOM | Q330A |
| 5 | CACGCGATGGTTCCGTTCTACGGGGCAGGCATGAACGCGGG | CACGCGATGGTTCCGTTCTACGGGGCAGGCATGAACGCGGGCTTTGAAGACTGTAGC | IF/IF HOM | Q330A |
| 6 | CACGCGATGGTTCCGTTCTACGGGGCAGGCATGAACGCGGG | CACGCGATGGTTCCGTTCTACGGGGCAGGCATGAACGCGGGCTTTGAAGACTGTAGC | IF/IF HOM | Q330A |
| 7 | CACGCGATGGTTCCGTTCTACGGGCCAGGCATGAACGCGGG | CACGCGATGGTTCCGTTCTACG--------------------TTGTAAGACTGTAGC | IF/FS | Q330P |
| 8 | CACGCGATGGTTCCGTTCTACGGGCCAGGCATGAACGCGGG | CACGCGATGGTTCCGTTCTACG--------------------TTGTAAGACTGTAGC | IF/FS | Q330P |
| 9 | CACGCGATGGTTCCGTTCTACGGGGCAGGCATGAACGCGGG | CACGCGATGGTTCCGTTCAACGGG----GCATGAACGCGGGCTTTGAAGACTGTAGC | IF/FS | Q330A |
| 10 | CACGCGATGGTTCCGTTCTACGGGGCAGGCATGAACGCGGG | CACGCGATGGTTCCGTTCAACGGG----GCATGAACGCGGGCTTTGAAGACTGTAGC | IF/FS | Q330A |
| 11 | CACGCGATGGTTCCGTTCTACGGGGCAGGCATGAACGCGGG | CACGCGATGGTTCCGTTCAACGGG----GCATGAACGCGGGCTTTGAAGACTGTAGC | IF/FS | Q330A |
| 12 | CACGCGATGGTTCCGTTCTACGGGGCAGGCATGAACGCGGG | CACGCGATGGTTCCGTTCATGT----GGGCATGAACGCGGGCTTTGAAGACTGTAGC | IF/FS | Q330A |
| 13 | CACGCGATGGTTCCGTTCTTGTGGGCAGGCATGAACGCGGG | CACGCGATGGTTCCGTTCAACGGG----GCATGAACGCGGGCTTTGAAGACTGTAGC | IF/FS | W328L; G329W; Q330A |
| 14 | CACGCGATGGTTCCGTTCTACGGGGCAGGCATGAACGCGGG | CACGCGATGGTTCCGTTCTACAACGCAAGGTTCTACGCGGGCTTTGAACATGAACGC | IF/FS | Q330A |

**Supplementary Table 13. Eye-phenotype proportions scored for *kh*$^{Rec+}$ vs *kh*$^{+R}$ allelic challenge.**

| Generations | Replicate Cages | | | | | | | | | | | |
|---|---|---|---|---|---|---|---|---|---|---|---|---|
| | A | | | B | | | C | | | D | | |
| | GFP$^{+}$ | GFP$^{–}$ | Total | GFP$^{+}$ | GFP$^{–}$ | Total | GFP$^{+}$ | GFP$^{–}$ | Total | GFP$^{+}$ | GFP$^{–}$ | Total |
| **G1** | 79.3% | 20.7% | 300 | 81.2% | 18.8% | 394 | 77.9% | 22.1% | 653 | 81.4% | 18.6% | 420 |
| **G2** | 80.7% | 19.3% | 373 | 85.9% | 14.1% | 396 | 82.3% | 17.7% | 362 | 87.9% | 12.1% | 390 |
| **G3** | 84.2% | 15.8% | 310 | 89.2% | 10.8% | 305 | 84.0% | 16.0% | 357 | 89.0% | 11.0% | 308 |
| **G4** | 86.8% | 13.2% | 318 | 90.0% | 10.0% | 359 | 85.7% | 14.3% | 357 | 89.6% | 10.4% | 326 |
| **G5** | 88.3% | 11.7% | 315 | 91.0% | 9.0% | 310 | 87.5% | 12.5% | 327 | 89.9% | 10.1% | 338 |
| **G6** | 89.4% | 10.6% | 320 | 95.7% | 4.3% | 304 | 88.9% | 11.1% | 341 | 90.8% | 9.2% | 336 |

Experiments were conducted in four replicate cages (A-D) each seeded with 200 individuals heterozygous for a copy of the Rec*kh* drive allele and a copy of the *kh* functional resistant allele (*kh*$^{Rec+}$/*kh*$^{+R}$) with a 1:1 sex ratio.
All mosquitos displayed WT black eye color.
